# CORTICAL AUDITORY PROCESSING FROM CHILDHOOD TO ADULTHOOD: ASSOCIATIONS WITH SPEECH UNDERSTANDING

**DOI:** 10.1101/2025.10.06.680107

**Authors:** Kumari Anshu, Didulani N. Dantanarayana, Shelly P. Godar, Ruth Y. Litovsky, Carlos R. Benítez-Barrera

## Abstract

The maturation of the auditory system is critical for the development of speech perception from childhood through early adulthood. However, the developmental trajectories and behavioral significance of cortical responses to speech sounds, particularly in relation to frequency specificity, remain poorly understood. Here, we presented low-frequency (/m/) and high-frequency (/s/) speech sounds to 60 typically developing individuals aged 5–24 years and recorded early cortical responses (P1 and N1) using electroencephalography. We also examined associations between these neural responses and speech understanding in quiet and in the presence of speech interferers. The developmental trajectories of P1 and N1 revealed distinct age- and stimulus-dependent patterns, including both linear and non-linear changes across development. These findings delineate frequency-specific maturational profiles within the cortical auditory system and identify potential neurophysiological markers of speech perception, providing a normative benchmark for assessing atypical auditory development.

## INTRODUCTION

The maturation of the human auditory system is a long and complex process, undergoing significant developmental changes from the embryonic period to early adulthood^1^. The peripheral auditory system is mature in infancy, but the central auditory system develops gradually. While the brainstem auditory pathway is fully developed by early childhood, the cortical auditory areas continue to mature well into late adolescence^2–4^. During the post-natal developmental period, auditory brain regions undergo structural and functional changes that are shaped by both inherent (genetic) and environmental (sensory) factors^5,6^.

The functional developmental trajectory of the central auditory system can be observed through neurophysiological responses to auditory stimuli recorded using electroencephalography (EEG) techniques^2,7^. Specifically, in awake, neurotypical individuals with normal hearing, passive listening to auditory stimuli elicits obligatory cortical auditory evoked potentials (CAEPs)^8,9^. These CAEPs result from the summation of temporally overlapping electrical activity at multiple neural sources and in adults, appear as a sequence of positive-negative-positive-negative deflections ∼50 to 300 ms post-stimulus onset, commonly referred to as the P1-N1-P2-N2 complex^10,11^. This multiphasic complex can be observed at the vertex or Cz (i.e., the center of the scalp) and Fz (midline frontal) electrodes, but its constituent peaks represent contributions from multiple distributed neural sources, and weight of these contributions depends on the position of the recording electrode^10,12^. The P1 is considered to arise from the lateral portion of Heschl’s gyrus, i.e. secondary auditory cortex, though it may also include activity from subcortical sources, the N1 has sources in the primary and secondary auditory cortices, and the P2 has a wider source distribution, including primary auditory cortex, auditory association cortices, and the reticular activating system^10,12–16^. Acoustic features of stimuli are known to influence the amplitudes and latencies of CAEP components. For example, differences in stimulus parameters such as frequency, level, inter-stimulus interval, rise time, spectro-temporal complexity, and speech vs. non-speech stimuli, all influence CAEP characteristics^17–19^. The latency and amplitude of CAEP peaks are also highly dependent upon electrode location, the integrity of the auditory pathway, and the organization of cortical auditory areas, including the primary and secondary auditory cortices^10,15,20^.

CAEPs, particularly the P1 and N1 responses, undergo age-related changes from infancy to adulthood that track developmental changes in the auditory cortex^4,10,11,21–23^. CAEP responses are present in newborns and young infants; however, the morphology, latency, amplitude, and scalp distribution of CAEPs during development differ greatly from adult responses^19,23,24^. Several maturational stages have been identified before CAEPs become fully mature in young adulthood^25–29^. In young children up to 5-6 years of age, a biphasic response is observed at frontal and central locations of the scalp (Fz and Cz electrodes)^10,23^. This response is characterized by a prominent, broad positive P1 peak (around 80-100ms post-stimulus onset) followed by a broad negative deflection (around 200 ms post-stimulus onset; referred to as N2). By 8-9 years of age, the N1 response first emerges as a small negative deflection in the broad positive P1 peak at electrodes Fz and Cz at a latency of around 100ms, the P2 becomes apparent, and the multiphasic response can be observed for the first time at these locations^10,28^. After this age, P1 gradually decreases in amplitude and latency, reflecting more mature physiology, as observed with faster and more efficient stimulus detection^22,30^. On the other hand, N1 increases in amplitude and decreases in latency with age until it becomes a fully mature discrete component in young adulthood, signifying fine-tuning of sensory representation of the auditory signal at the cortical level^3,22,26,27^. There are conflicting reports on age-related changes in P2 amplitudes and latency as P2 is greatly influenced by top-down processes, increasing response variability. Thus, P1 and N1 responses can serve as more reliable indicators of auditory system maturation than P2 and are well-suited for tracking developmental changes in cortical auditory processing^10,21,22,24,25,28,29^.

Despite extensive research on auditory cortical development using CAEPs, several areas remain underexplored. First, most seminal developmental studies used non-speech sounds, such as tones or clicks, to elicit CAEPs^10,21,22,25,28,31^, while only a few have examined maturational changes evoked by speech sounds^24,27,29,32^. Investigating the maturation of speech-evoked CAEPs in typically developing (TD) children offers a valuable tool for understanding the neurophysiological processes underlying the normative development of speech processing and provides insights into the neural encoding of speech in children at risk of neurodevelopmental delays^15,17,33^. Further, as speech-evoked CAEPs can be elicited during passive listening, this paradigm is suitable for individuals with neurodevelopmental differences for whom active task performance might be challenging.

Second, there is a paucity of studies specifically examining frequency-specific differences in the maturation of speech-evoked CAEPs. Behavioral studies show that sensitivity to high-frequency pure tones matures by as early as 6 months of age, whereas sensitivity to low-frequency pure tones matures later, i.e., by early school age (see review^34^). Discrimination of low-frequency pure tones continues to mature into the school years and is not yet fully developed by age 10, whereas high-frequency discrimination is already well developed by around age three^35^. These maturational differences may reflect underlying organizational principles observed in functional imaging studies which show that auditory regions are tonotopically organized. For example, in the primary auditory cortex, different frequencies activate distinct cortical areas, and distinct layers within a cortical column^36,37^. Given that the tonotopic organization of the auditory cortex continues to mature and refine with age^12,24,38^, CAEPs are likely to show different maturational trajectories based on the spectral characteristics of different speech stimuli. In the present study we were interested in capturing frequency-specific responses to speech-relevant stimuli; one approach for doing so is to use isolated phonemes, which are the smallest building blocks of speech sounds and exhibit dominant spectral energy in distinct frequency ranges^38,39^. Although some studies that investigate auditory maturation have examined CAEPs in response to phonemes, very few have directly compared responses to phonemes with distinct frequency profiles^18,19,38,40,41^.

Third, studies investigating speech-evoked CAEPs have generally employed a group design and focused on a narrow age range, targeting either infants, children, adolescents or adults^11,18,24,26,27,38^. Examination of CAEPs across a wide age range, from childhood to adulthood, would provide a comprehensive picture of the entire maturational process throughout development. This has the potential to aid in early assessment of atypical auditory developmental trajectory, for example, in children with hearing difficulties or in individuals with neuro-maturational delays who are at risk for communication challenges.

Fourth, the relationship between the maturation of speech-evoked CAEPs and speech perception abilities remains poorly understood. Speech perception depends on the neural encoding of multiple aspects of speech sounds, such as frequency, amplitude, and timing cues, which are reflected in CAEP components^17,42^. Thus, P1 and N1 obligatory responses are objective markers of a part of cortical processing that underlies complex skills such as understanding speech^15,43,44^. Speech perception, including understanding speech in quiet and in degraded conditions, develops throughout childhood and reaches adult-like proficiency by late childhood and late adolescence, respectively, depending on the specific task and type of stimulus used^45,46^. Thus, the protracted maturation of CAEPs is accompanied by improvements in auditory perceptual skills throughout the developmental period^2,4,15^. However, developmental improvements in speech perception are only partly due to changes in auditory processing throughout development. Maturation of the cognitive and language systems also contributes to improvement in speech perception when listening in complex acoustic environments (see reviews^34,47,48^). The present study addresses a timely need to better understand the association between speech recognition and CAEPs across development.

Very few maturational studies have specifically examined the relationship between speech perception and CAEPs in a population ranging from children to adults. Most previous studies have focused on specific age groups and yielded mixed findings (see reviews^17,42,49^). Differences in task stimuli, CAEP, and speech perception testing in quiet or real-world conditions (i.e., in the presence of background noise), sample size, age range of participants, and their language and cognitive development stage may contribute to the lack of consistent findings^43,50,51^. Most notably, there might be frequency-specific differences in the association between cortical maturation and the development of speech perception skills. High-frequency sounds are known to be critical for speech understanding, particularly in the presence of background noise^52–54^. Thus, CAEPs evoked by high-frequency speech sounds might better predict performance on speech perception tests in degraded conditions than CAEPs in response to low-frequency sounds. Understanding how frequency-specific auditory processes develop and relate to the development of speech perception abilities is crucial for designing interventions that ensure access to optimal auditory input, such as with ideally fitted hearing aids and/or cochlear implants.

The goal of the present study was to address these gaps in the literature by evaluating the auditory maturational process using CAEPs in response to low-frequency (/m/) and high-frequency (/s/) phonemes in neurotypical individuals with no known hearing difficulties, ranging in age from 5 to 24 years. This comprehensive characterization of age-related changes in CAEP can serve as a benchmark to assess auditory maturational deficits in individuals with hearing difficulties or neuro-maturational delays. We focused on the P1 and N1 peaks, which are reliable markers of auditory system maturation^24,28,29^. The following were the specific aims and hypotheses:

- To measure developmental trajectories of obligatory CAEP components in response to high-frequency (/s/) and low-frequency (/m/) sounds in neurotypical individuals (aged 5 to 24 years) with no known hearing difficulties. We hypothesized that younger children (5-8 year olds) would show immature biphasic response, which would gradually transition to a triphasic response. Further, that P1 amplitudes and latencies would decrease while N1 would exhibit a decrease in latency and an increase in amplitude from early childhood to adulthood. Additionally, consistent with the development of behavioral sensitivity, P1-N1 responses to high-frequency sounds were predicted to show signs of earlier maturation as compared to low-frequency sounds.
- To assess if CAEPs in response to /m/ and /s/ stimuli presented in quiet conditions would be associated with speech recognition skills in quiet and in the presence of speech interferers. We hypothesized that mature CAEP components (lower P1 and higher N1 amplitudes and shorter P1 and N1 latencies) would be associated with understanding of speech materials at lower sound levels, indicating better performance, in quiet and in the presence of background speech maskers. Further, we explored whether the association between CAEPs and speech recognition would differ for low- and high-frequency stimuli.

## METHODS

### Participants

Neurotypical children and adults (n=63) were recruited from the general Madison, WI area community. Children ranged in ages from 5-17 years (10.60 ± 3.35 years), and adults ranged in age from 18-24 years (20.49 ± 1.54 years). Criteria for participation included English as a primary spoken language and no known hearing loss, confirmed with pure tone audiometry (at or below 20 dB hearing levels in both ears at octave frequencies between 250 and 8000 Hz). Additional criteria for excluding participation were the presence of atypical neurological history, serious medical diagnoses, or developmental disabilities (e.g., autism spectrum disorder, intellectual disability, attention deficit disorder). Of these, three participants were excluded from the study because their pure-tone air conduction thresholds did not meet criteria. The final sample consisted of a total of 60 participants: 38 neurotypical children (16 females, 22 males) and 22 adults (19 females, 3 males). Each child participant was accompanied by a parent or legal guardian, and written informed consent and assent were obtained before participation. All adult participants provided written consent. Because the present study was part of a larger study involving multiple measures, participants completed testing across different days, ranging from 2-5 study visits. This study was approved by the Institutional Review Board at the University of Wisconsin-Madison.

### EEG measures

#### Stimuli

Auditory stimuli consisted of naturally produced /m/ and /s/ phonemes, created from speech samples ‘ama’ and ‘asa’, spoken by the same adult male talker and were recorded with a sampling frequency of 44.1 kHz. Each stimulus had a duration of 80 milliseconds (ms), including 15 ms of cosine squared ramps (Figure 1). The speech sounds /m/, / and /s/ have a spectral emphasis in the low-, and high-frequency regions (spectral centroid ∼200 Hz, and ∼6000 Hz, respectively).

**Figure 1:**
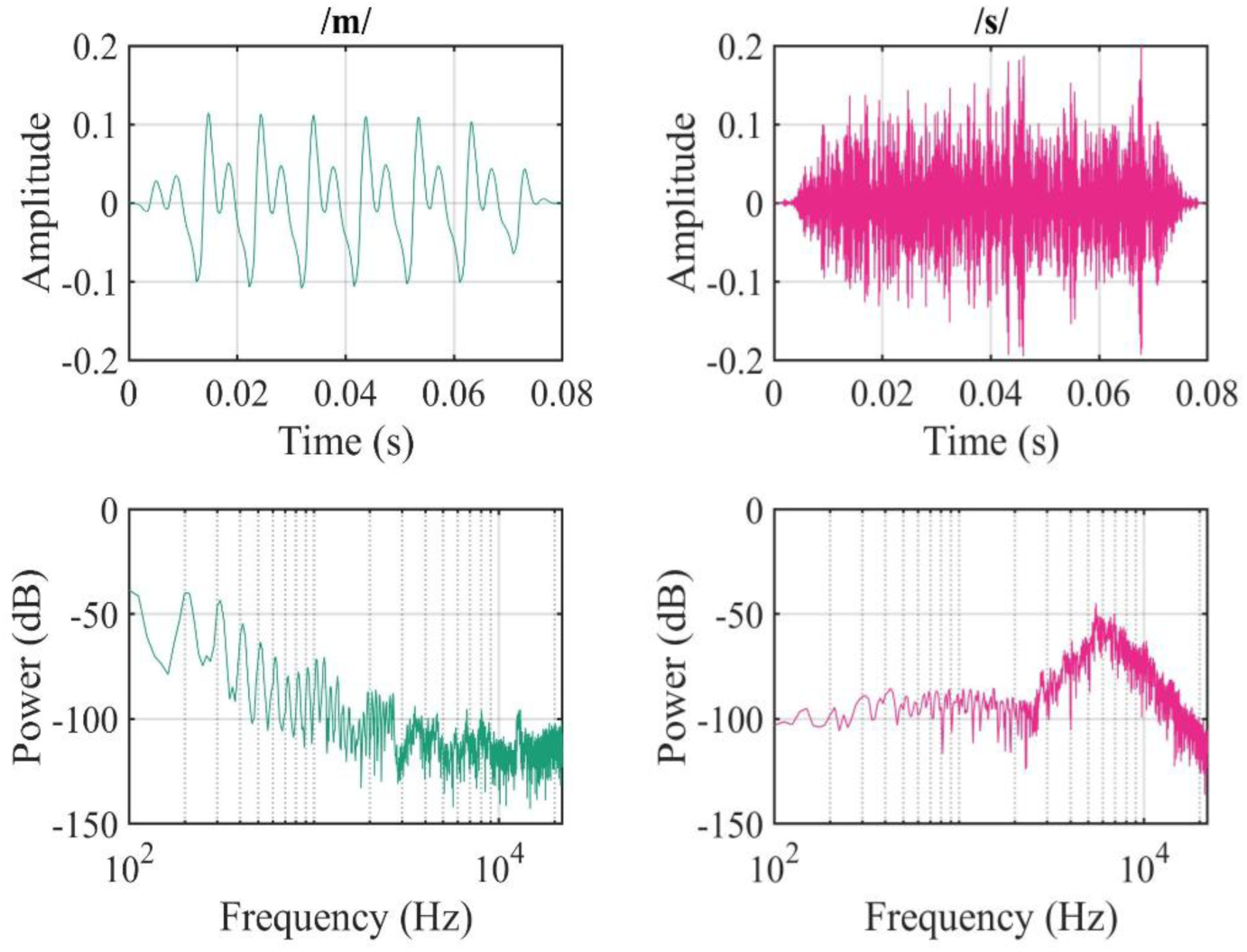
**Waveforms and power spectra of /m/ and /s/ stimuli.**

#### EEG recording

EEG data were acquired using a 128-channel Hydrocel Geodesic Sensor Net connected to a NetAmps 400 amplifier and Net Station 5.0 software (Magstim Inc, USA). The vertex channel (Cz) served as the reference during data collection. The impedances of all channels were kept at or below 50 kOhms and EEG signals were acquired at a sampling rate of 1000 Hz. All recordings were conducted in a standard IAC (Industrial Acoustics Company) sound-attenuated booth with an ambient temperature maintained at 68°F. Audio stimuli were presented using EPrime 3.0 software (Psychology Software Tools, USA). To ensure accuracy and precision of stimulus onset time in synchronization with EEG, the stimuli were routed through a StimTracker (Cedrus, San Pedro, CA, USA) that was connected to the earphones, and the EEG amplifier.

During the EEG session, participants passively listened to auditory stimuli while watching a silent video, chosen by them from a list of age-appropriate videos. Participants were instructed to ignore the test stimuli, focus on the video, and sit as still as possible. Stimuli were presented monaurally to one ear at a time at 65 dB SPL using ER-2 insert earphones (Etymotive Research). During the testing for one ear, the other ear was not occluded. The order of testing right or left ear condition was counterbalanced across participants. Both /m/ and /s/ phonemes were randomly presented (150 trials/phoneme), with an interstimulus interval (ISI) ranging from 1000 to 2000 ms. The primary tester continuously monitored the EEG data quality during acquisition. A second tester sat in the participant room to closely observe the participant for any major movements. Recordings were paused if there were significant movement artifacts. The testing was resumed once the participant was ready and able to sit still. Testing took around 7-8 minutes per ear. Prior to our EEG task, participants completed a resting state task where they were asked to sit still with their eyes open and then closed three times for 1 minute each. However, note that resting EEG data are not discussed here. The entire EEG session, including cap placement, impedance checking, data acquisition, and breaks between different conditions lasted on average 1.5 hrs.

#### EEG Data Preprocessing and Analysis

EEG preprocessing was performed offline using custom scripts in EEGLAB v2023.1^55^ within the MATLAB v2021b environment (Mathworks, Massachusetts, USA)^55^. Raw data were filtered using a low-pass Finite Impulse Response (FIR) filter at 30 Hz and a high-pass FIR filter at 0.5 Hz, re-referenced to the average, and downsampled to 512 Hz. The clean_rawdata plugin v2.91, implementing the Artifact Subspace Reconstruction (ASR) algorithm was applied to discard poor quality segments^56,57^. Channels were rejected if they were flat for over 5 s, showed excessive line-noise (z > 4) relative to the recording, or were not strongly correlated with neighbors (r < 0.80). For calibration, data were high-pass filtered (0.25–0.75 Hz transition band), and transient bursts were identified at 20 standard deviations. Artifact windows were rejected if over 25% of channels were contaminated and only extreme high-power segments were rejected. Missing channels were interpolated using spherical methods. The artifact free continuous data were then epoched from −100 to +500 ms relative to the onset of the auditory stimulus (/m/ and /s/ phonemes) and baseline-corrected using the pre-stimulus period (-100 to 0 ms). Only data from participants who had at least 50 clean trials for each phoneme in each ear condition were included in statistical analyses, in line with recommended thresholds^58,59^.

Epoched data were then further processed for CAEP analysis using ERPLAB plugin v10.04^60^ in EEGLAB. P1 and N1 amplitudes and latencies were extracted using automated peak detection with the *pop_geterpvalues* function within ERPLAB^60^. For all participants, the P1 and N1 peaks were defined as the most positive or negative deflection at Cz occurring within 30–110 ms and 70–150 ms, respectively. These time windows were selected based on prior auditory maturation studies and further verified through visual inspection of the grand-averaged waveforms for each subject^10,24,27,29^. If visual inspection revealed that the peak was outside the default window, the ERPLAB measurement tool was used to manually adjust the window and identify the peak amplitude and latency^11,26^. Notably, the algorithm identified a peak only if the value was higher or lower (for P1 and N1, respectively) than the preceding and subsequent three consecutive data points. For both P1 and N1, the baseline-to-peak amplitude and latency from stimulus onset values were extracted for each participant for further analyses. Although CAEPs were recorded monoaurally, no significant between-ear differences were observed in amplitude and latencies. To streamline the analysis, we averaged the P1 and N1 values across both ears and used them for subsequent analyses. After screening for an adequate number of artifact-free trials, EEG data from 59 participants were included in the final analysis.

### Speech Understanding Task

#### Stimuli

A set of 25 bisyllabic words (spondees) spoken by a male talker was used as target stimuli^61^. The interferer stimuli consisted of two talker speech interferers generated by superimposing two recordings of a single male talker speaking different sentences from the Harvard IEEE corpus^46,61^. The interferer stimuli did not have any silent gaps or pauses.

#### Task Procedure

The Children’s Realistic Index for Speech Perception (CRISP), a closed-set four-alternative forced-choice (4-AFC) task, was used to measure speech understanding in quiet and three different conditions with background interferers^61^. Testing was completed in a single-walled sound-treated IAC booth. Target stimuli were always presented from a loudspeaker in front of the listener (0° azimuth). Each auditory target word had a corresponding visual icon. On each trial, four of the target icons were presented on a touchscreen monitor, such that one of the icons matched the auditory stimulus. A trial began with an alerting phrase (“Ready?”) preceding the target word, after which the four icons were displayed. Participants were asked to select the icon that matched the perceived word. Following each incorrect response, feedback was provided.

To represent different real-world listening conditions, in which interferers might come from different directions, we implemented three different conditions with interfering speech: Front (interferers co-located with the target); Right (interferers at 90° azimuth to the right of the participant); Left (interferers at 90° azimuth to the left of the participant). Loudspeakers were located at 1.2 m from the center of the participant’s head. In total, there were four listening conditions (Quiet, Front, Right, and Left), and testing was repeated twice for each condition, in randomized order. During each test run, the target stimulus level was varied adaptively with a starting level of 60 dB SPL. Initially, following each correct response, the target level decreased by 8 dB, and following the first incorrect response, a modified 3-down/1-up rule was used (see ref.^61^ for detailed adaptive tracking rules). The level of interfering stimuli was fixed at 55 dB SPL. This adaptive protocol has been used extensively in previous speech understanding studies in adults and children aged 3 years and older^46,61–63^. To ensure familiarity with target words, participants completed a familiarization session before testing began, to confirm their ability to associate each word with its corresponding icon. The protocol specifies that if a given target word was unfamiliar to a participant, that word was removed from the list of options for that participant. While a minimum familiarity with 15 target words was required to qualify, all participants successfully identified the full set of 25 words, allowing the entire set to be used in testing.

Speech reception thresholds (SRTs) were determined using a logistic function fitted to data using the MATLAB “psignifit” toolbox (version 4)^64^, as described by Wichmann and Hill^65^, and similar to our prior analysis approach for these type of data^46,62,63,66,67^. The SRT corresponded to the 79.4% correct point on the psychometric function, with the guessing rate (γ) fixed at 0.25 and the lapse rate (λ) capped at 0.06. Thresholds from all runs within a condition were averaged to produce a single value per participant.

### Statistical Analysis

Our first aim was to characterize age-related changes in CAEPs (P1 and N1 amplitudes and latencies) in response to each phoneme (/m/ and /s/). Since CAEPs might exhibit non-linear changes with age^10,11^, we modeled the relationship of each CAEP measure with stimulus type and age using both generalized additive models (GAMs) and linear mixed-effects models (LMMs) with linear, polynomial, and logarithmic terms for age. Model selection was based on criteria such as lower Akaike Information Criterion (AIC) and higher conditional R² (for LMMs) or higher deviance explained (for GAMs). GAM models that included separate smooth terms for /m/ and /s/ consistently explained the most deviance and are reported here, while quadratic models which had the lowest BICs are reported in supplementary results.

To complement these models with age as a continuous variable, we also conducted group-based comparisons by dividing participants into four developmental age groups: Younger children (5-8 years old, n=12), Older children (9-12 years old, n=14), Adolescents (13-17 years old, n=11) and Adults (18-24 years old, n=22). These age groups were created based on prior literature documenting key maturational changes in P1 and N1 waveform characteristics^10,24,28,29^. Around 7-8 years of age, N1 begins to emerge with long ISIs (1000 ms and more), and by 9-10 years, most children exhibit reliable N1 peaks. By ∼12 years of age, a stable triphasic CAEP waveform is typically observed. From age 13 onwards, CAEPs continue to mature gradually, and by adulthood, N1 is considered fully developed. The age groups were therefore selected to capture these meaningful stages of auditory cortical maturation. CAEPs were also analyzed using LMMs with fixed effects for age group, stimulus type, and their interaction. Random intercepts for subjects were included, and Holm correction was applied to all post hoc pairwise comparisons.

To address our second aim of exploring the relationship between CAEPs and behavioral speech understanding (SRTs), we first computed zero-order correlations between CAEP measures, age, and SRTs in each listening condition (Quiet, Front, Right, Left). We also examined bivariate correlations between CAEPs and SRTs correcting for multiple comparisons. To account for developmental effects, we employed LMMs with SRTs as the dependent variable and CAEP measure, listening condition, and their interaction as fixed effects. Age was included as a covariate, and each model included a random intercept for subjects. Because CAEP measures are not independent, we ran separate LMMs for each measure. Separate models were also run for /m/ and /s/ stimuli, as including both phonemes within a single model resulted in convergence issues. Given the exploratory nature of this aim, we did not correct for multiple testing across all models. All predictors were mean-centered, and model residuals were examined for any violation of normality and homoscedasticity assumptions. All statistical analyses were conducted using R software (version 4.2.2; R Core Team 2022).

## RESULTS

### Descriptive Developmental Changes in CAEPs Morphology and Scalp Topography

A qualitative evaluation of CAEP morphology and scalp topography of P1 and N1 components across age groups was carried out. Supplementary Table 1 provides the average number of accepted trials per phoneme for each group. Grand average waveforms at the Cz electrode for each age group are shown in Figure 2A and 2B. Developmental changes in the CAEP waveform morphology can be observed for both the /m/ and /s/ phonemes. In adults, the CAEP response featured a prominent N1 peak around 100 ms, preceded by a smaller P1 peak at approximately 50 ms. In contrast, generally children’s waveforms exhibited a larger P1 and a smaller N1 than adults. Scalp topography maps at the grand average peaks for each group are shown in Figure 2C and 2D.

**Figure 2:**
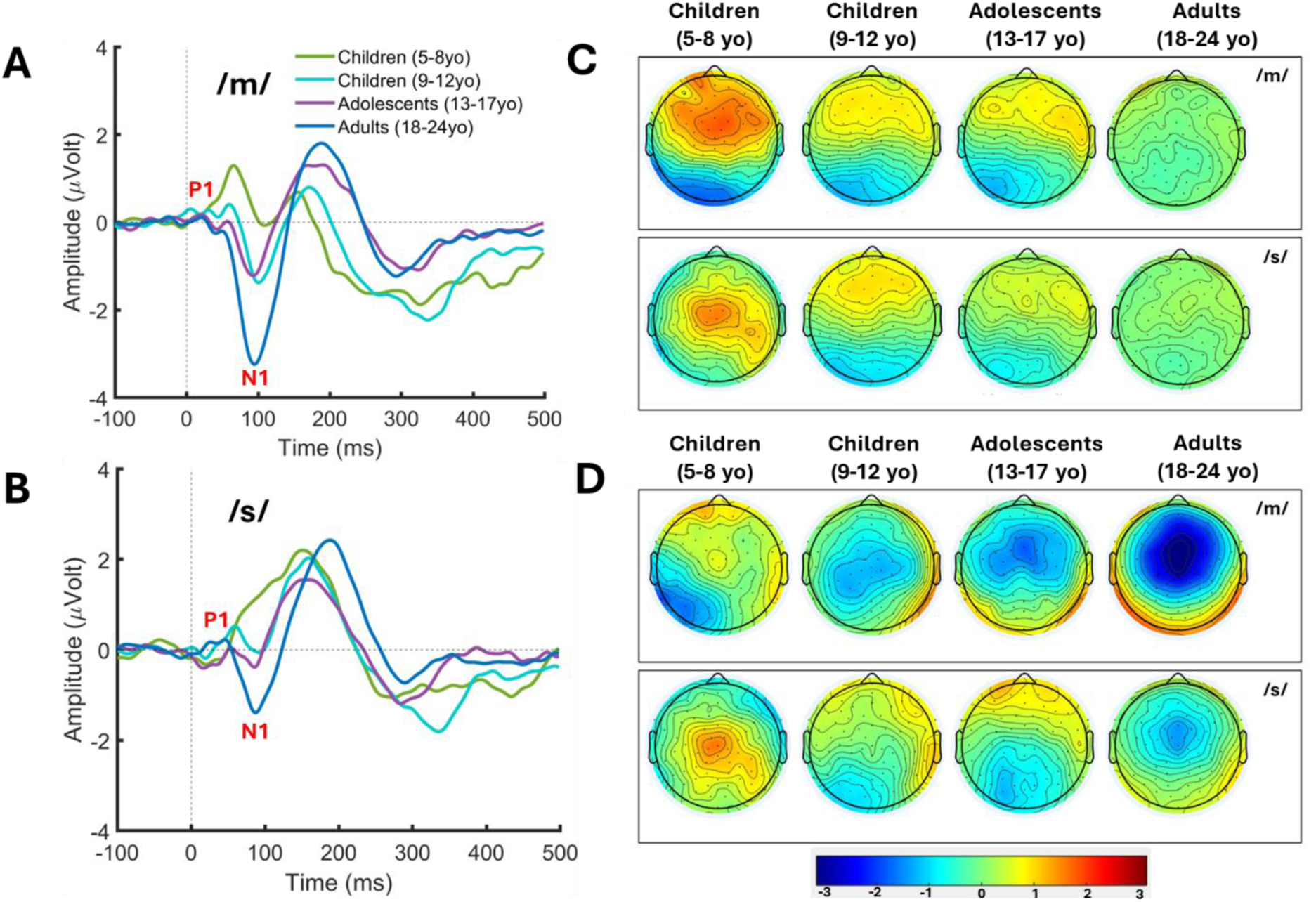
Grand average waveforms and scalp topographies. Grand average waveforms at electrode Cz show age-related changes in P1 and N1 morphology across four age groups in response to /m/ and /s/ stimuli (A, B). Scalp topography plots at the grand average P1 and N1 peak latencies illustrate developmental changes in response distribution and magnitude for each age group and stimulus (C-D). Warmer colors indicate positive voltage; cooler colors indicate negative voltage.

In children aged 5-8, CAEP waveforms for /s/ stimuli showed a broad positive response followed by a small dip, which evolved into the N1 peak in older children (9-12 years). The N1 peak became more prominent in adolescence (13-17 years old) and reached full maturity in adults. In contrast, for /m/ stimuli, children aged 5-8 years did not exhibit a biphasic response. Instead, they showed a triphasic response with a distinct N1 peak, albeit with a much smaller amplitude than that observed in older children and adults. Like the /s/ stimuli, the N1 amplitude in response to /m/ stimuli increased progressively across age groups. The topography plots show that for ages 5-8 years, P1 was stronger in frontocentral regions for /m/ and central regions for /s/, while N1 was small for /m/ and absent for /s/. As age increased, the P1 response weakened. In contrast, the N1 response, which showed a parietal distribution in the youngest children, gradually shifted to a frontocentral distribution with age and became prominent and centrally focused in adults.

### Developmental Changes in CAEP Amplitude and Latency Across Different Age Groups

To characterize age-related maturational changes in P1 and N1 responses to /m/ and /s/ stimuli, the amplitude and latency of P1 and N1 peaks were compared across different age groups of children and adults (Figure 3).

**Figure 3:**
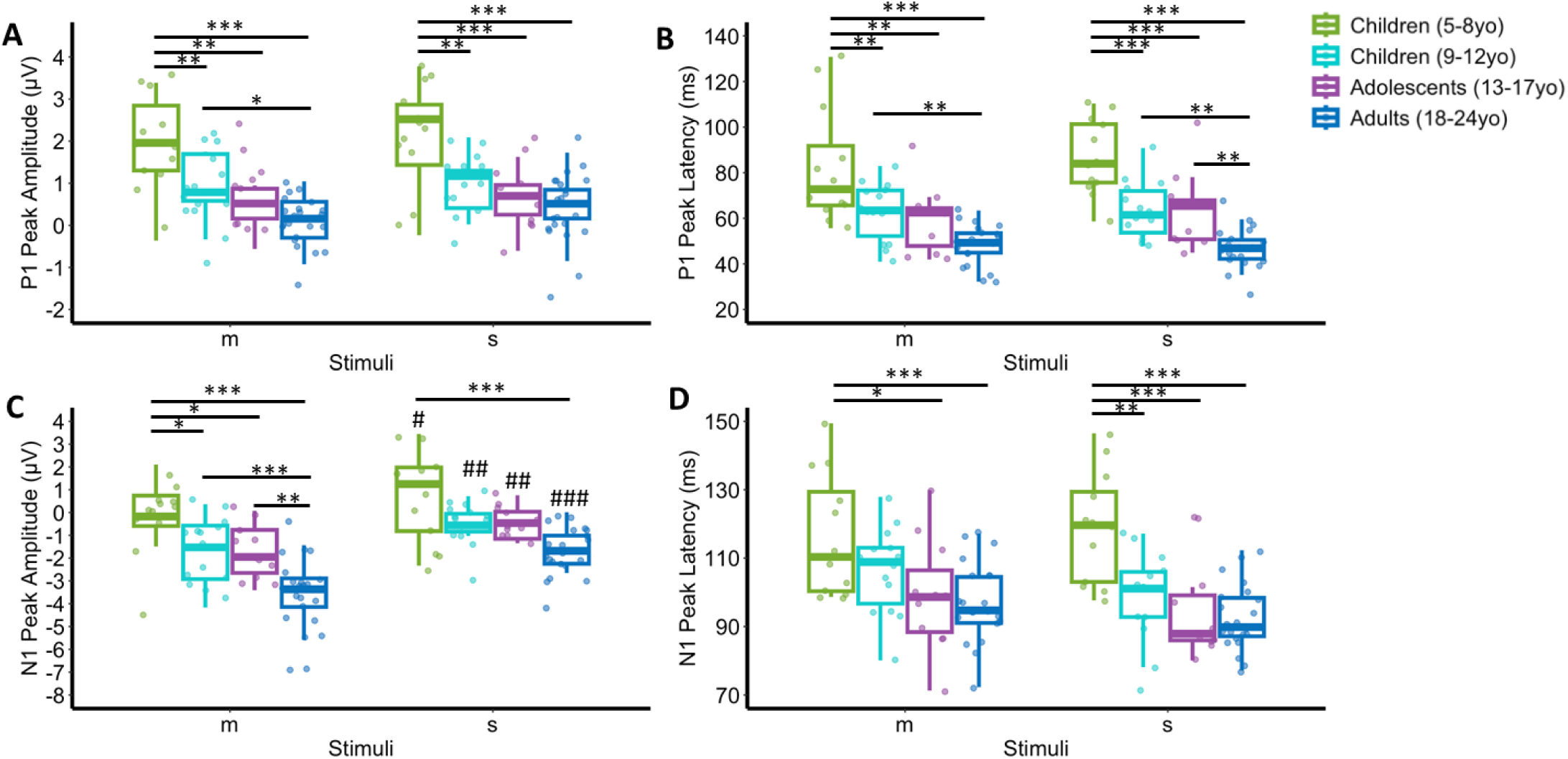
Age related differences in CAEPs for /m/ and /s/ stimuli. Box plots show the distribution of P1 peak amplitude (A), P1 peak latency (B), and N1 peak amplitude (C) and N1 peak latency (D) across four age groups for both /m/ and /s/ stimuli. Individual data points are overlaid on each box plot. Symbols: Differences between age groups (*), Differences between stimuli (#), The number of symbols indicate p values, for example, *p<0.05; **p<0.01; ***p<0.001.

#### P1 amplitude and latency

Significant main effects of age group were observed (Figure 3A-3B; amplitude: F(3,55) = 16.815, p<0.001; latency: F(3,55) = 24.156, p<0.001). There were no significant main effects of stimulus, nor a significant age group x stimulus interaction, indicating that age-related changes in P1 amplitude and latency were not different for /m/ and /s/ stimuli. Within each age-group, no significant differences in P1 amplitude or P1 latency were found between /m/ and /s/ stimuli. Between-group comparisons for /m/ stimuli revealed that children aged 5-8 years had significantly higher P1 amplitudes than those aged 9-12 years, adolescents and adults (all p≤0.002). Additionally, children aged 9-12 years had significantly higher P1 amplitude than adults (p=0.044), whereas no significant differences were found between adolescents and adults. For /s/ stimuli, children aged 5-8 years had significantly higher P1 amplitudes compared to children aged 9-12 years, adolescents and adults (all p≤0.001). There were no other group differences in P1 amplitude for /s/ stimuli. Between-group comparisons for P1 latency showed significantly longer P1 latency in children aged 5-8 years compared to all other groups for both /m/ stimuli (all p≤0.003) and /s/ stimuli (all p<0.001). Among older age groups, children aged 9-12 years had longer latencies than adults for both stimuli (/m/: p=0.009; /s/: p=0.003), whereas adolescents had longer P1 latency than adults only for /s/ stimuli (p=0.004).

#### N1 amplitude and latency (Figure 3C-3D)

Significant main effects of age group were found (amplitude: F(3,55) = 16.039, p<0.001; latency: F(3,55) = 10.466, p<0.001), indicating age-dependent changes in both measures. A significant main effect of stimulus was found for amplitude (F(1,55) = 57.263, p<0.001) but not for latency (F(1,55) = 3.392, p=0.071), suggesting that N1 amplitude, but not latency, varied by stimulus-type. There were no significant interactions between age group x stimulus for N1 amplitude or latency. Within-group comparisons revealed that N1 amplitude was significantly larger for /m/ stimuli than /s/ stimuli in all groups (all p≤0.02). A longer N1 latency for /m/ compared to /s/ stimuli was observed only in children aged 9-12 years (p=0.044). Between-group comparisons revealed that for /m/ stimuli children aged 5-8 years had significantly lower N1 amplitudes compared to children aged 9-12 years, adolescents and adults (all p<0.02), while for /s/ stimuli, they showed lower amplitudes only compared to adults (p<0.001). Children aged 9-12 years, and adolescents had lower N1 amplitudes compared to adults for /m/ (p<0.001, and p=0.001, respectively), but not for /s/ stimuli. For N1 latency, between-group comparisons revealed that children aged 5-8 years had longer latencies as compared to adolescents (p=0.016) and adults (p<0.001) for /m/ stimuli and compared to all the other age groups for /s/ stimuli (all p≤0.001). No other significant group differences in N1 latency were observed for either stimulus.

### Associations Between Age and P1 and N1 Amplitude and Latency

Across all our GAM models, the explained deviance ranged from 77% to 84%, indicating strong model fits (Figure 4). Subject-level random effects were significant for all CAEP measures (effective degrees of freedom [edf] > 30, p<0.001), highlighting substantial individual variability in auditory maturation.

**Figure 4:**
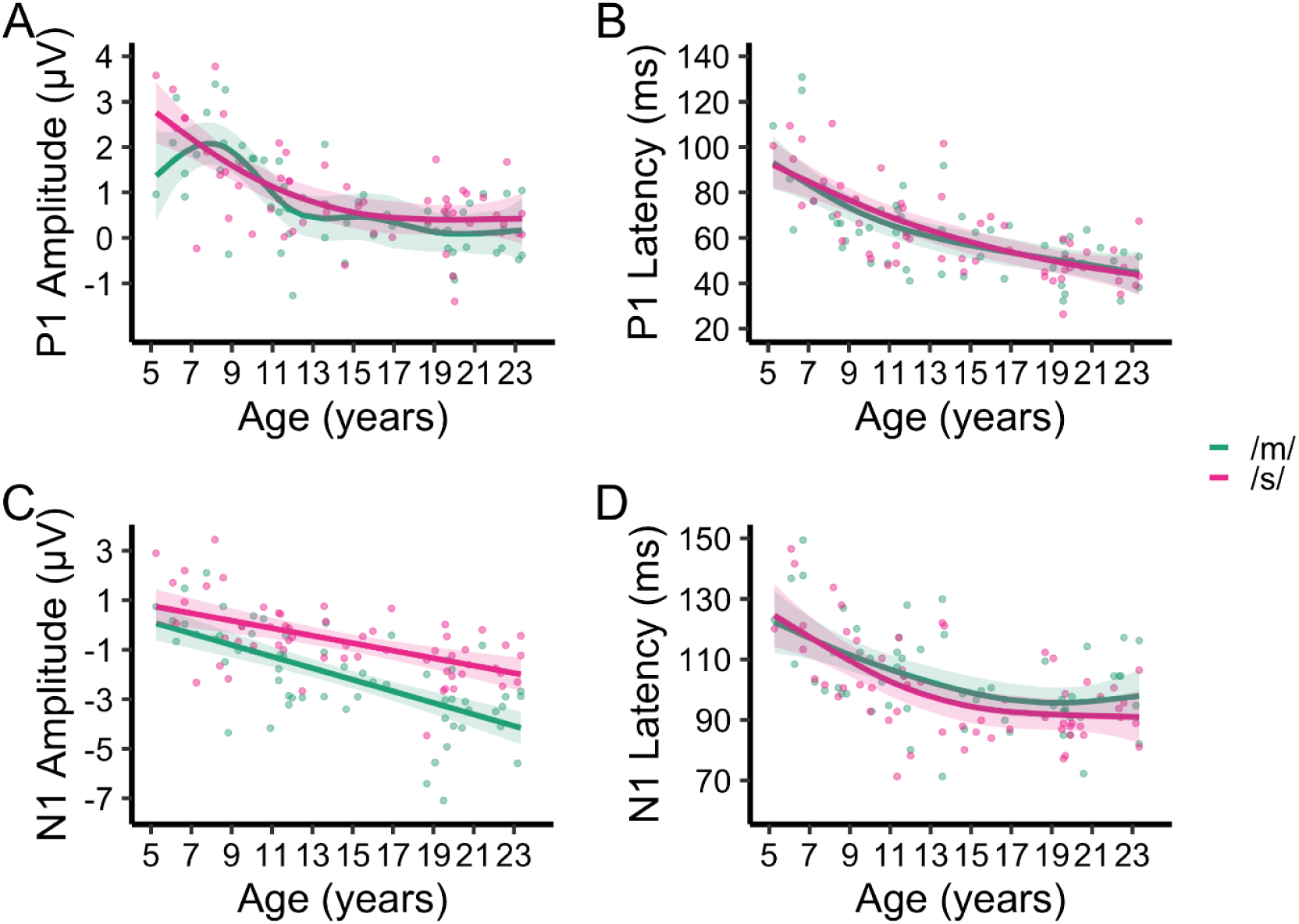
Relationships between Age and CAEPs for /m/ and /s/ stimuli. GAM model fits showing developmental trajectories of P1 and N1 amplitude and latency across age for /m/ (green) and /s/ (magenta) stimuli. Shaded regions represent 95% confidence intervals. Panels illustrate age-related changes for: (A) P1 amplitude, (B) P1 latency, (C) N1 amplitude, and (D) N1 latency.

#### P1 amplitude and latency

Age-related changes in P1 amplitude were highly non-linear, especially for /m/ stimuli (edf = 5.58, F = 8.34, p<0.001), suggesting a complex maturational trajectory. For /s/ stimuli, the relationship was also non-linear, but less complex (edf = 2.48, F = 15.6, p<0.001). Notably, stimulus-specific differences were found in the youngest individuals, pointing to early developmental differences in how /m/ and /s/ are processed. In contrast, the relationship of P1 latency with age was significant but smoother for both /m/ and /s/ stimuli (/m/: edf = 2.47, F = 18.8, p<0.001; /s/: edf = 1.95, F = 28.3, p<0.001). There were no significant differences between stimuli, suggesting a shared developmental pattern for P1 latency across phonemes.

#### N1 amplitude and latency

Unlike P1, N1 amplitude increased linearly with age (edf = 1) for both stimuli with strong effects (/m/: edf = 1, F = 52.5, p<0.001; /s/: edf = 1, F = 22.0, p<0.001). To further explore potential stimulus-specific differences in maturation, a linear regression model revealed a significant age × stimulus interaction (β = 0.0824, p=0.009), indicating steeper age-related increases in N1 amplitude for /m/ than for /s/. This suggests that N1 responses to low-frequency phonemes continue to strengthen across development, while responses to high-frequency phonemes follow a shallower trajectory and remain less prominent even in adulthood. Although a quadratic model had a slightly lower BIC, the age-squared term was not significant, so a linear model was retained for interpretability and consistency across CAEP measures. Details of model comparisons and results from the linear and quadratic models are summarized in Supplementary results. N1 latency showed modest but significant non-linear relationships with age for both stimuli (/m/: edf = 2.23, F = 9.12, p<0.001; /s/: edf = 2.23, F = 13.0, p<0.001). However, there were no stimulus-specific differences in the relationship with age, suggesting a largely shared developmental pattern for N1 latency.

### Relationship between CAEPs and Speech Reception Thresholds (SRTs)

The initial approach was to check zero-order correlations between all CAEP measures (P1 and N1 amplitudes and latencies), SRTs in each listening condition, and age. Note that all SRT and CAEP measures were significantly correlated with age. Supplementary Figure 1 illustrates the linear relationships between pairs of variables and their Spearman correlation coefficients and p-values, as well as histograms of each variable. Pairwise Spearman correlations between SRTs and CAEPs were then examined, following correction for multiple comparisons (Supplementary Table 2). Except for the relationship between N1 and P1 amplitudes with SRTs in Quiet, SRTs in all listening conditions were significantly correlated with CAEPs. However, these bivariate relationships between P1 and N1 measures and SRTs could be driven by age since both these measures are highly correlated with age and show developmental changes.

Results from linear mixed effect models examining the relationship between SRTs and CAEPs across listening conditions, including age as a covariate, are reported here. Figure 5 shows the relationships for P1 and N1 amplitudes with SRTs in Quiet and in the presence of co-located (Front) and separated (Right and Left) interferers for /m/ and /s/ stimuli. Figure 6 shows the corresponding results for P1 and N1 latencies.

**Figure 5:**
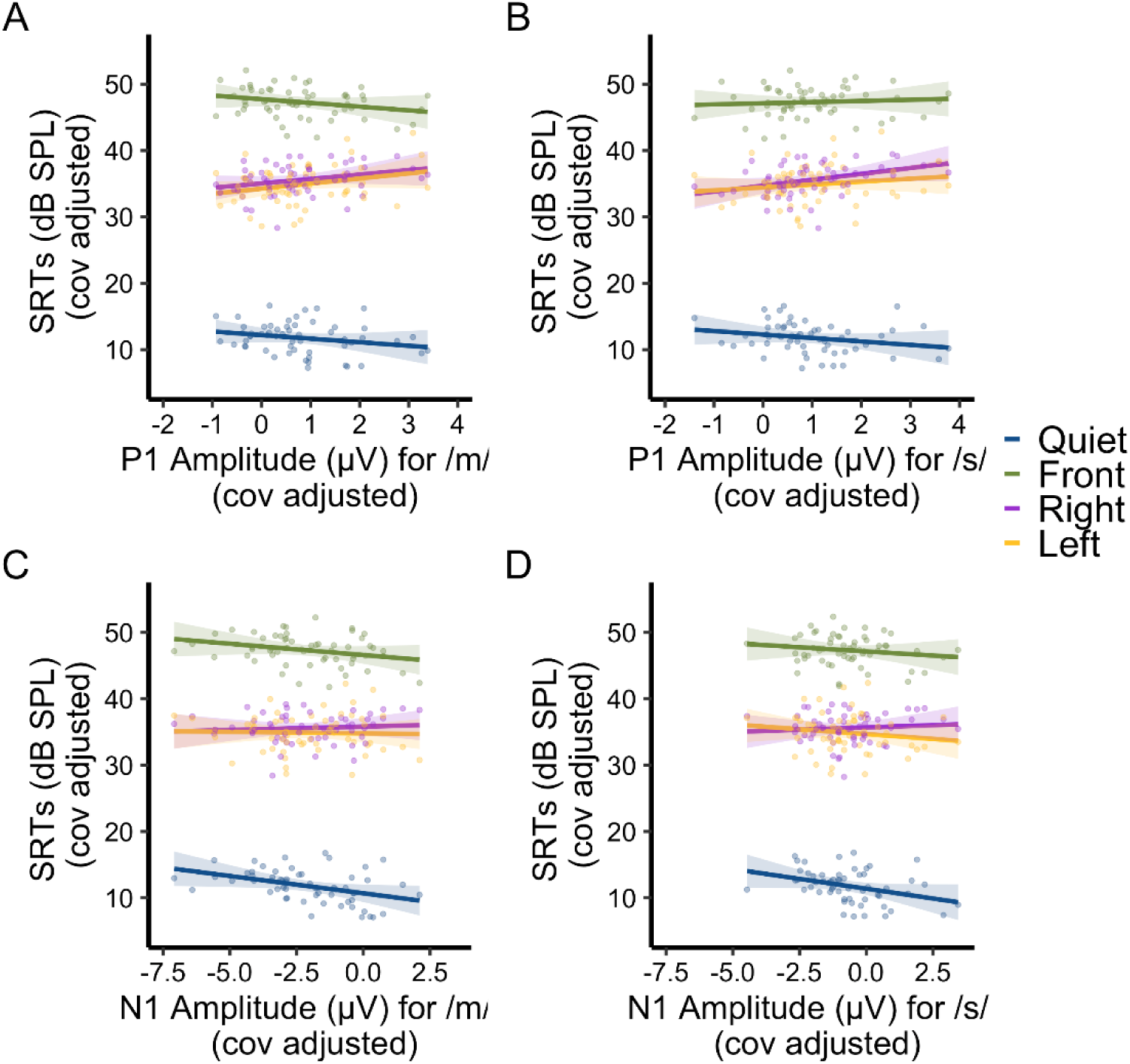
Relationships between CAEP amplitudes and SRTs for /m/ and /s/ stimuli. Scatterplots and regression lines show the associations between SRTs (in dB SPL) and P1 (A, B) and N1 (C, D) peak amplitudes for /m/ and /s/ stimuli, across four listening conditions, based on the location (if any) of the interferers. Shaded regions represent 95% confidence intervals. Age was included as a covariate in the linear mixed-effects models.

**Figure 6:**
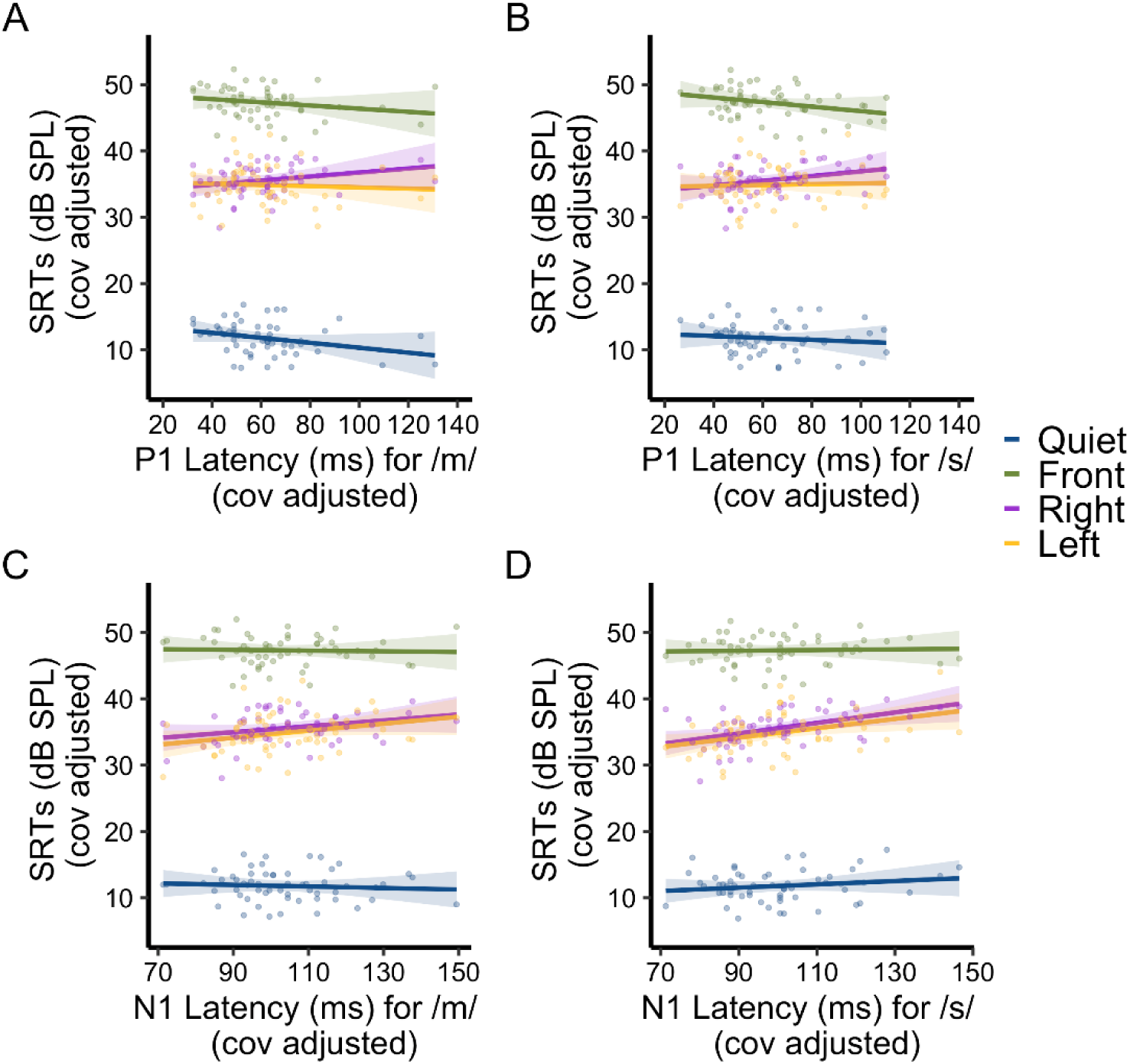
Relationships between CAEP latencies and SRTs for /m/ and /s/ stimuli. Scatterplots and regression lines show the associations between SRTs and P1 (A, B) and N1 (C, D) peak latencies for /m/ and /s/ stimuli across the four listening conditions, based on the location (if any) of the interferers. Shaded regions represent 95% confidence intervals. Age was included as a covariate in the linear mixed-effects models.

#### P1 amplitude

For /m/ stimuli, P1 amplitude was not a significant predictor of SRTs (F(1,55) = 0.048, p=0.827), but there was a significant main effect of condition (F(3,167) = 1153.628, p<0.001) and a P1 amplitude x condition interaction (F(3,167) = 4.629, p=0.004). The positive relationships of P1 amplitudes and SRTs in Right and Left conditions were significantly different than the Quiet condition (p=0.013 and p=0.007 respectively). Similarly, for /s/ stimuli, there was no main effect of P1 amplitude (F(1,55) = 0.491, p=0.486), but there was a significant main effect of condition (F(3,167) = 922.951, p<0.001) and P1 amplitude x condition interaction (F(3,167) = 3.037, p=0.031). The relationship of SRTs with P1 amplitude in the Right and Left conditions for /s/ were significantly more positive than the Quiet condition (p=0.004 and p=0.047 respectively).

#### N1 amplitude

There were no significant main effects of N1 amplitude for either stimulus (/m/: F(1,55) = 0.673, p=0.315; /s/: F(1,55) = 1.096, p=0.299). There was a main effect of listening condition for both stimuli (/m/: F(3,167) = 870.195, p<0.001; /s/: F(3,167) = 1505.052, p<0.001). There were no significant interactions between N1 amplitude x condition for either stimulus (/m/: F(3,167) = 2.447, p=0.065; /s/: F(3,167) = 1.541, p=0.206).

#### P1 latency

For /m/ stimuli, P1 latency did not significantly predict SRTs (F(1,55) = 0.256, p=0.615), but there was a main effect of condition (F(3,167) = 154.289, p<0.001) and a P1 latency x condition interaction (F(3,167) = 2.662, p=0.049). The relationship of SRTs with P1 latency in the Right condition for /m/ was significantly more positive than in the Quiet condition (p=0.008). A similar pattern was found for /s/ stimuli. P1 latency was not a significant predictor of SRTs (F(1,55) = 0.006, p=0.937), but there was a main effect of condition (F(3,167) = 168.960, p<0.001) and a significant P1 latency x condition interaction (F(3,167) = 2.791, p=0.042). The relationship of P1 latency and SRTs in the Right condition for /s/ was again more positive than the relationship in Quiet (p=0.049).

#### N1 latency

For /m/ stimuli, N1 latency was not a significant predictor of SRTs (F(1,55) = 0.879, p=0.353). There was a significant main effect of condition (F(3,167) = 35.855, p<0.001), but no significant N1 latency x condition interaction (F(3,167) = 2.124, p=0.099). For /s/ stimuli N1 latency significantly predicted SRTs (F(1,55) = 4.423, p=0.040), and there was a main effect of condition (F(3,167) = 47.673, p<0.001). The interaction between N1 latency x condition was marginal (F(3,167) = 2.647, p=0.051). Post hoc comparisons revealed significant associations between N1 latency and SRTs in the Right (p=0.006) and Left (p=0.015) conditions, but not in the Front (p=0.856) and Quiet (p=0.383) conditions.

## DISCUSSION

The present study investigated central auditory system maturation using CAEPs in response to low frequency /m/ and high frequency /s/ speech stimuli in individuals 5 to 24 years of age. By examining maturational changes in P1 and N1 amplitude and latency from childhood to adulthood, and their associations with speech reception thresholds (SRTs), we aimed to characterize typical auditory system maturation. In addition, these findings are intended to provide a benchmark for analysis of maturation of the auditory system in special populations such as children with hearing loss or with neurodevelopmental delays. We found distinct age- and stimulus-dependent changes in morphology, amplitude, and latency measures of CAEPs. The N1 peak latency was the only neural marker that significantly predicted behavioral performance on the speech understanding task (in the Right and Left conditions, i.e., when interferers were spatially separated from the target), even after accounting for the age of the particpant.

### Changes in CAEP morphology and topography across development

A critical feature of CAEP maturation is the change in morphology of the waveforms with age. In accordance with earlier studies, for both /m/ and /s/ stimuli, the grand average waveform at Cz for children 13-17 years old tested here had a triphasic morphology with a clear negative peak (N1) at around 100 ms that became more prominent in adults^10,28^. Furthermore, the grand average waveform for /s/ stimuli for youngest age group of children (5-8 years old) exhibited a classic biphasic response as per the literature, with an emerging notch suggestive of an N1 peak in the next oldest age group (9-12 years old). For /m/ stimuli, 5-8 year old children exhibited a larger N1 dip compared to /s/, but the N1 morphology remained immature compared to 9-12 year old children who had a clear triphasic morphology. There are at least two explanations for the differences between responses to /m/ and /s/ among 5-8-year-olds. First, in general, /m/ stimuli elicited a robust N1 response with much larger N1 peak amplitude as compared to /s/ stimuli for all participants. Second, there was large individual variability in CAEP waveform shapes among children 5-8 years old. For example, two 8-year-old children had very prominent N1 peaks for /m/ stimuli that overlapped with the range of N1 peak amplitudes for adults and were clearly different from the rest of the children in this age group, who showed no clear N1 peak (See supplementary figure 2 showing individual waveforms for all participants in this age range). Variance in auditory maturation is a hallmark of many studies, suggesting that auditory sensitivity and processing of complex signals mature at different rates amongst children of the same age^10,15,21,28^. Future studies with larger sample sizes in younger children (< 8 years old) are needed to continue to improve our understanding of the implications of these findings.

Scalp topography for the P1 response showed a fronto-central distribution for both /m/ and /s/ stimuli among children ages 5-8 years, shifting anteriorly toward a more frontal distribution primarily in the 9-12 year age group, with minimal subsequent changes in adolescents and adults. Prior studies, typically using limited electrodes, reported a predominantly fronto-central P1 distribution stable from 6-10 years of age^10,21,23,24^, with anterior shifts occurring gradually between 10-18 years of age^22,26,27^. Notably, our high-density 128-channel EEG recordings revealed an earlier and more pronounced shift among children 9-12 years of age. These differences might reflect improved spatial resolution provided by higher-density EEG, as well as methodological variations including stimulus characteristics, age groupings, analysis windows, and referencing schemes.

Clear developmental shifts were also observed in N1 scalp topography. Among children aged 5-8 years, the N1 response to /m/ was maximal over parietal regions, shifting gradually to fronto-central areas in children 9-12 years of age and adolescents up to 17 years of age, and becoming focused fronto-central in young adults (18-24 years). In contrast, for /s/, the N1 distribution was not present in children aged 5-8 years, emerged in the parietal region in middle childhood, gained prominence in adolescence, reaching a fronto-central pattern in adulthood. Prior reports on N1 developmental topography are somewhat mixed. Several studies have reported a shift from a parietal to a fronto-central distribution between ages 6-15 years^28,68,69^, while others describe a more lateral and widespread frontal distribution in 10-14 year-olds that becomes increasingly focused and midline in adolescents and adults^27,70^. Our results are broadly consistent with the view that N1 becomes more fronto-central and spatially focused with age with some stimulus-specific differences. For example, we find that for /m/ stimuli these shifts begin early, particularly during the 9-12 year age range, whereas for /s/ stimuli, our N1 topography findings suggest a later emergence and a more gradual transition to a fronto-central distribution by adulthood. There might also be differences in spatial resolution, stimulus characteristics, age group definitions, or the analytic time windows used, which could contribute to the slightly different patterns observed in the present study.

### Changes in P1 and N1 amplitude and latency across development

Consistent with our hypotheses, we observed robust age-related changes in the amplitude and latency of auditory P1 and N1 components in response to speech stimuli. Generalized additive models (GAM) revealed strong non-linear associations between age and P1 amplitude and latency. For both measures, there was a sharp decline in early childhood that gradually plateaued through adolescence into adulthood. P1 amplitude and latency did not differ significantly across stimuli, except for a modest divergence in amplitude in children younger than 7 years of age. Analyses by age group showed that children had significantly larger P1 amplitudes and longer latencies compared to adolescents and adults. These findings are consistent with previous studies showing P1 as a prominent component in early childhood that diminishes with maturation^10,24,29,32^.

For N1, amplitude increased linearly with age, but the rate of change differed by stimulus: /m/ showed a steeper increase compared to the more gradual rise observed for /s/, leading to larger between-stimulus differences in N1 amplitudes with age. These findings align with our hypothesis that cortical processing of high-frequency speech sounds matures earlier than that of low-frequency sounds, with each following distinct maturational trajectories. For N1 latency, we observed an overall age-related decrease, with some non-linearities in younger children, but no stimulus-specific effects. These results align with prior studies showing that N1, typically absent in early childhood, gradually develops with the maturation of thalamo-cortical and cortical pathways and becomes fully mature in adults^10,15,24,28^.

Notably, across all age groups, N1 amplitude was significantly greater for low-frequency /m/ than for high-frequency /s/ stimuli, consistent with previous adult studies using the same speech stimuli^38^ and with non-speech low- and high-frequency tonal stimuli^24,71^. The /m/ and /s/ stimuli might engage different neural populations due to the tonotopic organization of the auditory cortex, where low-frequency sounds activate more superficial regions yielding stronger scalp responses^37,38,72^. However, frequency alone does not fully explain the findings, as not all low-frequency phonemes elicit stronger N1 responses than the /s/ phoneme^38^. Instead, N1 amplitude differences between /m/ and /s/ may reflect the distinct spectrotemporal profiles of the stimuli: /m/ is a low-frequency, periodic, voiced nasal sound, while /s/ is a high-frequency, aperiodic, voiceless fricative^19,38,73,74^. Thus, both spectral-temporal complexity and phonetic features are likely to be involved in shaping N1 responses.

No other study has compared CAEP responses to /m/ and /s/ stimuli among children, limiting cross-study comparisons. Wunderlich and colleagues reported no differences in N1 amplitudes in response to low- and high-frequency tones among children aged 4.5 to 6.5 years^24^. In our study, the children in the youngest group (aged 5-8 years) were slightly older, and the significant continuous age x stimulus interaction revealed increasing age-linked N1 amplitude differences between stimuli, suggesting that the 5-8 year age range may reflect a transitional period during which N1 amplitude differences between low- and high-frequency stimuli begin to become more distinct.

In contrast to the stimulus-dependent differences in N1 amplitudes, we did not find such differences in P1 amplitude in any age group in our study. This is consistent with the lack of P1 amplitude differences for these stimuli among adults reported in a prior study^38^. On the other hand, studies examining infants comparing /m/ with a different high-frequency phoneme (/t/) reported lower P1 amplitudes for /m/ as compared to /t/^18,19^. P1 is prominent in infancy and in children under about six years of age, and its amplitude decreases with age thereafter^10,23,29^. We also observed this pattern in our study, where our continuous age analyses revealed early (before age 7) stimulus-linked differences in P1 amplitude which were absent in subsequent years. The absence of stimulus-dependent P1 amplitude differences found in the youngest age group may reflect the inclusion of slightly older children (until age 8) within that group. Overall, smaller P1 amplitudes for both stimuli may explain the lack of significant stimulus specific amplitude differences in both our study and the prior adult study. Another possible explanation is that the neural generator for P1, located in the lateral portion of the Heschl’s gyrus, is more sensitive to temporal complexity than to spectral complexity, and may underlie the reduced sensitivity of P1 amplitude to stimulus-specific spectral differences^38,75^.

Taken together, our findings indicate that P1 reflects general auditory detection processes, while N1 is a more refined marker of auditory cortical processing that is sensitive to the spectral complexity of speech stimuli. Our findings of distinct developmental trajectories of N1 amplitude in response to /m/ and /s/ stimuli suggest that different rates of development in frequency-specific neural processing are reflected in N1 amplitude, potentially involving different sets of underlying neural generators with different rates of maturation.

### Associations Between CAEPs and Speech Reception Thresholds

The second aim of this study was to examine how maturational differences in CAEPs, specifically P1 and N1 amplitude and latency, elicited by the onset of low- and high-frequency speech sounds presented in quiet, are related to word recognition performance in both quiet and noisy listening conditions. We hypothesized that more mature CAEP responses (i.e., lower P1 amplitude, higher N1 amplitude, and shorter P1 and N1 latencies) would be associated with better (lower) SRTs^52–54^.

This hypothesis was partially supported by our findings. N1 latency for /s/ stimuli emerged as a significant predictor of SRTs, with shorter latencies associated with better speech understanding performance. Longer (delayed) N1 latencies were significantly associated with higher (poorer) SRTs in the Right and Left spatial masker conditions, but not in the Quiet or Front (co-located masker) conditions. These results suggest that delayed low-level auditory processing of high-frequency speech sounds are associated with poorer ability to extract speech information in the presence of background maskers that are spatially separated from the target speech. This neural-behavioral association might reflect an underlying developmental process regarding how children utilize spatial cues to extract speech information from their environment. Spatial cues generally lead to improved performance related to the Front condition, but performance in the presence of spatial cues also introduces between-subject variability in SRTs, which results in detection of meaningful associations with neural timing. Notably, shorter N1 latency reflects faster neural processing. Prior work suggests that individuals with faster neural timing may have better access to temporal and spectral cues important for speech perception, particularly in conditions where speech and noise are spatially separated^76^.

Although N1 latency for /s/ stimuli did show a significant relationship with SRTs, the overall variance explained was modest, likely due to the influence of multiple factors. First, CAEPs were recorded in quiet, whereas the speech perception task involved active word understanding in both quiet and noise conditions. Prior research has shown that onset CAEPs recorded in quiet are only weakly related to speech perception in participants with typical hearing^76–78^, whereas stronger associations have been observed when CAEPs are recorded in the presence of noise^43,49^. Second, onset CAEPs were recorded under passive listening conditions and only reflect the early neural encoding of spectro-temporal properties of speech stimuli but behavioral speech understanding, particularly in complex environments, relies on a broader perceptual system that also includes attention, memory, and other cognitive processes. Studies have found that associations between CAEPs and speech perception are stronger when active task paradigms are used, which engage advanced auditory processing with attention or top-down modulation^79,80^. Finally, the stimuli used to elicit CAEPs were isolated phonemes, while the behavioral task required recognition of bisyllabic words. Prior work suggests that neural–behavioral correlations are better when the neural stimuli closely resemble the speech material used in the behavioral test^81^.

That said, the finding that N1 latency to /s/ is significantly associated with SRTs on a word recognition task among the typically developing children and adults in our study is very promising and suggests clinical applications. This is especially valuable for populations where reliable behavioral speech testing is difficult, such as very young children, individuals with atypical hearing (e.g., auditory neuropathy), or individuals with cognitive or language challenges, where passively recorded CAEPs might offer an objective marker of cortical auditory processing capacity relevant to speech perception^17,18,82^.

In summary, our findings suggest that N1 latency for /s/ stimuli may serve as a meaningful neural correlate of speech understanding performance in complex listening environments. While CAEP measures may not fully predict speech-in-noise skillss in all listening conditions, they provide insight into the maturational readiness of auditory cortical systems that support successful speech processing.

### Strengths, Limitations and Future directions

The present study had several strengths. Our participants spanned a wide and continuous age-range of typical development, enabling us to capture early non-linearities in auditory maturation tracked by changes in CAEPs. The use of high-density EEG offered the opportunity to examine traditional CAEP measures at the vertex channel, and in addition, visualize more fine-grained topographical changes in response to stimuli across age. Our task enabled assessment of speech perception in four different listening conditions allowing us to capture neuro-behavioral associations and avoid floor and ceiling effects in task performance. The differential maturation patterns observed for /m/ and /s/ stimuli highlight the importance of frequency-specific speech processing in understanding auditory development. Overall, the study characterized typical auditory maturation, revealed associations with speech understanding, demonstrated the existence of substantial individual variability in development, and charted overall trajectories for the P1 and N1 CAEP components. The robust age-related changes in CAEP measures suggest that these components can serve as objective markers for assessing auditory maturation and identifying developmental delays in clinical populations.

There are several limitations to consider. First, the study might have been underpowered to detect weaker associations between CAEP components and speech understanding in the Quiet and Front (co-located) masker conditions. Second, while speech understanding was measured in both quiet and noise, CAEPs were only assessed in quiet, limiting direct comparison across measures. Finally, although this study included only typically developing participants, a passive task was used to ensure the paradigm remains applicable for populations that are more difficult to test such as individuals with developmental delays. However, in typically developing individuals, the addition of active tasks might have yielded stronger associations with behavioral performance.

The study underscores the need to explore additional factors that contribute to associations between auditory cortical processing and behavioral measures of speech understanding in challenging environments. Two potential directions are considered relevant for future studies. First, the task used here for behavioral measures of speech understanding was relatively easy, the goal being to benchmark performance across a wide age range. A task that involves more complex speech processing such as understanding sentences would add depth to the topic and provide insight into associations with cortical processing of speech during development. Second, future studies could incorporate speech-in-noise stimuli for cortical potential recordings and employ task designs that include both passive and active paradigms. This would enable the assessment of both lower-level sensory encoding and higher-level speech-related cortical processing involving top-down modulation, helping to disentangle the respective contributions of different levels of auditory processing and cognitive factors to behavioral performance. Additionally, longitudinal designs would provide a more nuanced understanding of individual developmental trajectories and their variability.

## CONCLUSION

Our findings demonstrate significant developmental changes in CAEP waveform morphology, characterize the developmental trajectories of P1 and N1 amplitudes and latencies for both low-frequency /m/ and high-frequency /s/ phonemes, and reveal frequency-specific differences in auditory maturation. N1 latency for /s/ stimuli appears to be a meaningful neural correlate of speech understanding performance in noisy listening conditions. This study provides a comprehensive characterization of auditory cortical maturation in response to high- and low-frequency phonemes across a wide developmental age range. The findings highlight the dynamic nature of cortical auditory system maturation and its relevance for speech perception, offering a benchmark for future research on both typical and atypical auditory development.

## ACKNOWLEDGEMENTS

We thank all the participants and their families for their time and commitment to this study. This work was supported by the National Institutes of Health–National Institute on Deafness and Other Communication Disorders (NIH–NIDCD) under grants R01 DC003083 (Litovsky) and R01 DC019511 (MPIs: Litovsky, Hartley, and Alexander). Additional support was provided by a core grant to the Waisman Center from the National Institute of Child Health and Human Development (P50 HD105353). We are grateful to Stephanie Sellner, Aditi Gargeshwari, Mohammad Maarefvand, Molly Osborn, Elizabeth Neubauer, Amanda Lackner, and other members of the Binaural Hearing & Speech Laboratory for their help with data collection and study coordination.

