## supplementary for "CORTICAL AUDITORY PROCESSING FROM CHILDHOOD TO ADULTHOOD: ASSOCIATIONS WITH SPEECH UNDERSTANDING"

#### Supplementary Results:

#### Model comparisons and results for N1 amplitude and latency

To evaluate the developmental trajectories of N1 amplitude and latency, we compared multiple models including linear, quadratic, logarithmic, and generalized additive models (GAMs), with both main effects and interaction terms for stimulus type. Model fit was assessed using explained variance (conditional $R^2$ or deviance explained), Akaike Information Criterion (AIC), and Bayesian Information Criterion (BIC).

For N1 amplitude, the GAM interaction model provided the best overall fit ($R^2 = 0.85$, AIC = 346.82), outperforming the quadratic interaction model, which had lower BIC (406.25) but weaker fit ($R^2 = 0.76$).

For N1 latency, the quadratic interaction model yielded the lowest AIC (897.60) and BIC (919.76), whereas the GAM interaction model explained more variance ($R^2 = 0.77$) at the cost of higher BIC. Given this trade-off, the GAM model is reported in the main text for consistency, and the quadratic model is documented here for reference.

##### Linear and Quadratic Model Results

##### N1 Amplitude

- Linear Model: A significant age × stimulus interaction was observed ($\beta = 0.0824$, $p=0.009$), suggesting that the rate of maturation differed between the two phonemes. The N1 response to /m/ showed a steeper increase in amplitude with age than /s/, which exhibited a more gradual developmental trajectory.
- Quadratic Model: The main effect of the linear age term was highly significant ($\beta = -13.8$, $p<0.001$), while the quadratic term trended toward significance ($\beta = 3.43$, $p=0.070$). The age × stimulus interaction (linear term) remained significant ($\beta = 4.86$, $p=0.010$), reinforcing stimulus-specific differences in developmental trajectories.

##### N1 Latency

- Linear Model: Age was a significant predictor of N1 latency ($\beta = -1.33$, $p<0.001$), indicating shorter latencies with increasing age. However, the interaction between age and stimulus was not significant, suggesting similar maturational slopes for /m/ and /s/.
- Quadratic Model: Both the linear ($\beta = -78.3$, $p<0.001$) and quadratic ($\beta = 54.0$, $p=0.0035$) age terms were significant, indicating a modest non-linear trajectory for N1 latency across development. The main effect of stimulus was also significant ($\beta = -3.58$, $p=0.047$), but interaction terms were not significant, consistent with a largely shared developmental profile across phonemes.

#### Supplementary Tables:

##### Supplementary Table 1: Average number of clean EEG trials (mean± SD) across participants in each ear for /m/, and /s/ stimuli

| Ear | Stimulus | TDC1 | TDC2 | TDT | TDA |
|---|---|---|---|---|---|
| LE | /m/ | 117.8 ± 22.5 | 123.9 ± 14.3 | 129.8 ± 10.2 | 108.3 ± 24 |
| LE | /s/ | 117.1 ± 21.3 | 123.4 ± 13.8 | 128.5 ± 12.9 | 107.3 ± 22.5 |
| RE | /m/ | 115.6 ± 20.3 | 118.1 ± 14.3 | 132.1 ± 7.3 | 106 ± 25.2 |
| RE | /s/ | 116 ± 19.6 | 115.9 ± 15.9 | 130.8 ± 6.2 | 105.1 ± 25.1 |

LE: Left Ear
RE: Right Ear
TDC1: Children (5-8yo)
TDC2: Children (9-12yo)
TDT: Adolescents (13-17yo)
TDA: Adults (18-24yo)

**Supplementary Table 2: Correlations between SRTs and CAEPs.**

Spearman correlations between speech reception thresholds in each listening condition and P1 and N1 amplitudes and latencies for /m/ and /s/ stimuli sorted by FDR corrected p value.

| SRTs | CAEPs | Spearman Correlations | p | pFDR |
|---|---|---|---|---|
| Right | s_p1l | 0.54 | 1.25e-05 | 0.0002 |
| Right | m_p1l | 0.54 | 1.42e-05 | 0.0002 |
| Right | s_p1a | 0.53 | 3.07e-05 | 0.0003 |
| Right | s_n1l | 0.51 | 5.10e-05 | 0.0004 |
| Front | s_p1a | 0.50 | 9.55e-05 | 0.0005 |
| Right | m_p1a | 0.50 | 1.03e-04 | 0.0005 |
| Right | m_n1a | 0.48 | 1.69e-04 | 0.0007 |
| Left | s_n1l | 0.48 | 1.82e-04 | 0.0007 |
| Left | m_n1l | 0.44 | 7.03e-04 | 0.0025 |
| Left | m_p1a | 0.44 | 8.01e-04 | 0.0026 |
| Quiet | s_p1l | 0.42 | 1.12e-03 | 0.0030 |
| Front | s_n1l | 0.42 | 1.26e-03 | 0.0030 |
| Left | s_p1l | 0.42 | 1.27e-03 | 0.0030 |
| Right | s_n1a | 0.42 | 1.32e-03 | 0.0030 |
| Quiet | s_n1l | 0.41 | 1.47e-03 | 0.0031 |
| Left | s_p1a | 0.40 | 2.25e-03 | 0.0045 |
| Front | s_p1l | 0.38 | 3.29e-03 | 0.0062 |
| Right | m_n1l | 0.38 | 3.84e-03 | 0.0068 |

| SRTs | CAEPs | Spearman Correlations | p | pFDR |
|---|---|---|---|---|
| Front | m_n1a | 0.37 | 4.93e-03 | 0.0083 |
| Left | m_n1a | 0.36 | 5.83e-03 | 0.0093 |
| Front | s_n1a | 0.35 | 7.45e-03 | 0.0114 |
| Left | m_p1l | 0.34 | 8.61e-03 | 0.0125 |
| Front | m_p1l | 0.33 | 1.24e-02 | 0.0173 |
| Quiet | m_n1l | 0.33 | 1.35e-02 | 0.0180 |
| Front | m_p1a | 0.32 | 1.67e-02 | 0.0214 |
| Quiet | m_p1l | 0.31 | 1.81e-02 | 0.0223 |
| Front | m_n1l | 0.30 | 2.50e-02 | 0.0296 |
| Left | s_n1a | 0.29 | 2.73e-02 | 0.0312 |
| Quiet | m_p1a | 0.25 | 5.71e-02 | 0.0630 |
| Quiet | m_n1a | 0.23 | 9.07e-02 | 0.0967 |
| Quiet | s_n1a | 0.22 | 9.89e-02 | 0.1021 |
| Quiet | s_p1a | 0.16 | 2.32e-01 | 0.2320 |

### Supplementary Figures:

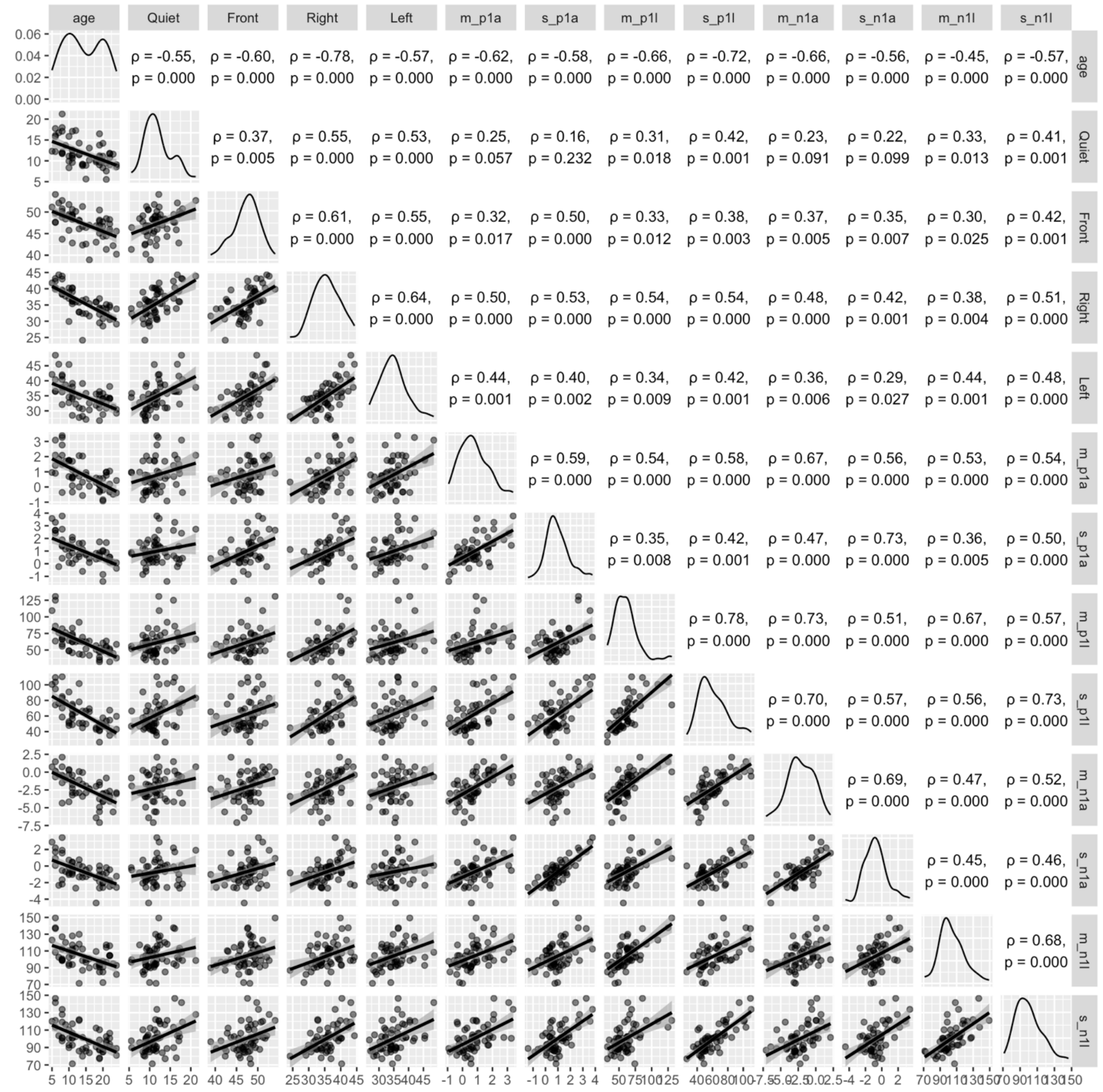


**Supplementary Figure 1: Zero-order correlations between Age, SRTs and CAEPs.** Diagonal elements indicate distribution of each variable. Scatterplots below the diagonal show pairwise-relationships between variables named on the row and column, and the corresponding intersections above the diagonals list the Spearman correlations and p values.

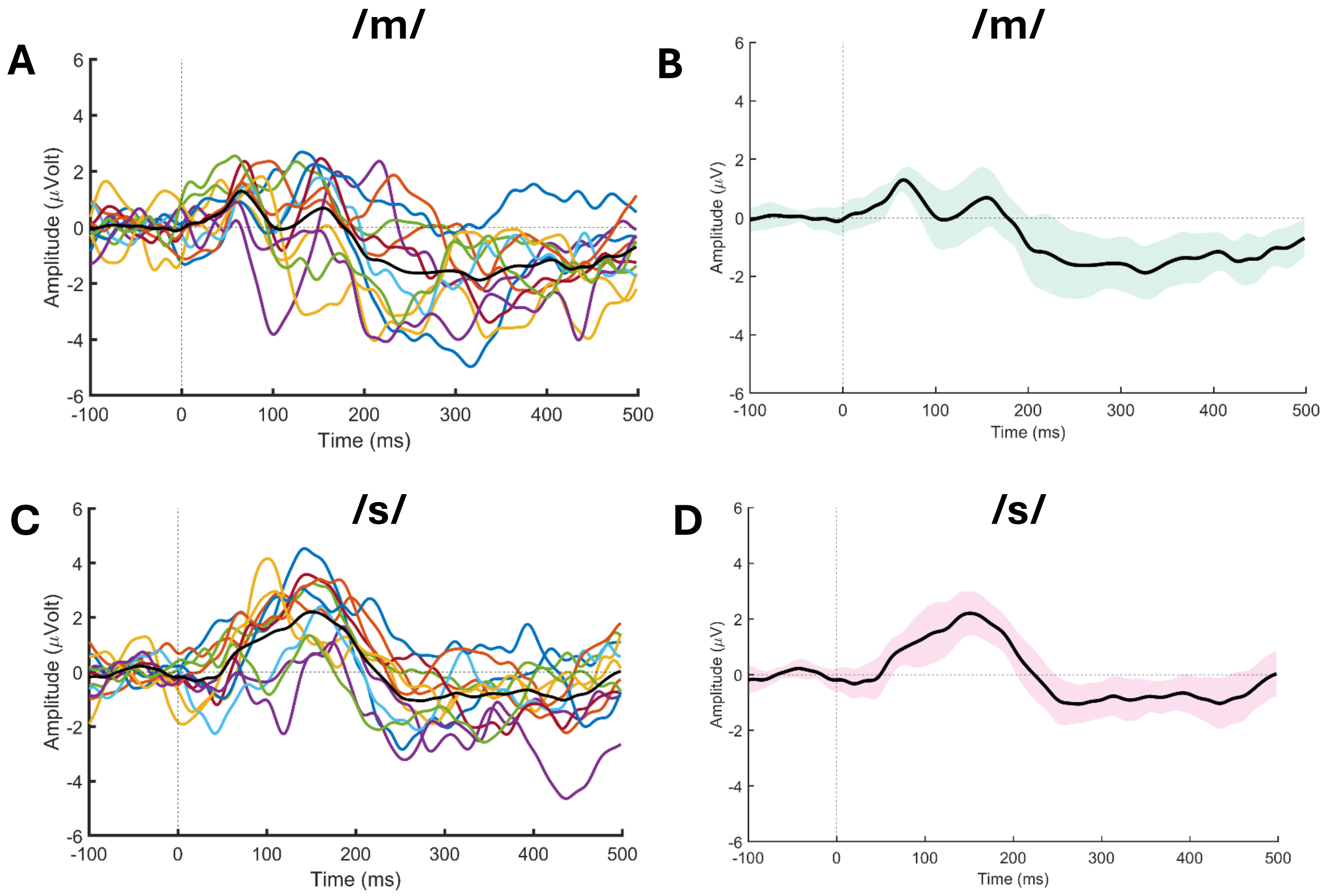


**Supplementary Figure 2. Individual and grand average waveforms for children aged 5-8 years.** Panels A and C show individual subject waveforms (colored lines) along with the group mean (black line) for the /m/ and /s/ stimuli, respectively. Panels B and D show the grand average waveforms with shaded 95% confidence intervals for the /m/ and /s/ stimuli.
